# Cohesin can remain associated with chromosomes during DNA replication

**DOI:** 10.1101/124107

**Authors:** James D. P. Rhodes, Judith H. I. Haarhuis, Jonathan B. Grimm, Benjamin D. Rowland, Luke D. Lavis, Kim A. Nasmyth

**Affiliations:** Department of Biochemistry, Oxford University, South Parks Road, Oxford, OX1 3QU, UK; Department of Cell Biology, the Netherlands Cancer Institute, Plesmanlaan 121, 1066 CX Amsterdam, the Netherlands; Janelia Research Campus, Howard Hughes Medical Institute, Ashburn, Virginia 20147, USA

## Abstract

To ensure disjunction to opposite poles during anaphase, sister chromatids must be held together following DNA replication. This is mediated by cohesin, which is thought to entrap sister DNAs inside a tripartite ring composed of its Smc and kleisin (Scc1) subunits. How such structures are created during S phase is poorly understood, in particular whether they are derived from complexes that had entrapped DNAs prior to replication. To address this, we used selective photobleaching to determine whether cohesin associated with chromatin in G1 persists in situ after replication. We used unlabelled HaloTag ligand following fluorescent labelling to block newly synthesised Halo-tagged Scc1 protein from incorporating fluorescent dye (pulse-chase or pcFRAP). In cells whose cohesin turnover is inactivated by deletion of *WAPL*, Scc1 remains associated with chromatin throughout S phase. These findings suggest that cohesion might be generated by cohesin that is already bound to unreplicated DNA.

## Introduction

The equal distribution of genetic material at cell division requires attachment of sister kinetochores to microtubules emanating from opposite sides of the cell, a process that depends on cohesion between sister chromatids. Sister chromatid cohesion is mediated by the cohesin complex, the core of which is a tripartite ring created by the binding of N- and C-terminal domains of a kleisin subunit Scc1 (Rad21) to the ATPase domains at the apices of a V-shaped Smc1/3 heterodimer (Gruber et al., 2003; Haering et al., 2002). Cohesion is thought to depend on entrapment of sister DNAs inside these tripartite rings while topologically associating domains (TADs) have been postulated to arise through the extrusion of loops of chromatin fibres through these rings, a hypothesis known as loop extrusion (Alipour and Marko, 2012; Fudenberg et al., 2016; Nasmyth, 2001; Sanborn et al., 2015).

The interaction between cohesin and DNA is regulated by the binding to Scc1 of three related regulatory subunits composed of HEAT repeats, namely Scc2 (Nipbl), Scc3 (SA1 and SA2), and Pds5, known collectively as HAWKs (Heat repeat containing proteins Associated With Kleisins) (Wells et al., 2017). Loading of cohesin onto chromosomes begins in telophase (Sumara et al., 2000) and depends on a complex of Scc2 and Scc4 (Mau2) (Ciosk et al., 2000). From this point until the onset of DNA replication, the dynamics of cohesin’s association with chromatin is determined by the rate of loading catalysed by Scc2 and the rate of release catalysed by Wapl (Kueng et al., 2006). During G1, these two processes create a steady-state where about 50% of cohesin is associated with chromatin with a residence time of about 20 minutes (Gerlich et al., 2006; Hansen et al., 2016).

Crucially, establishment of sister chromatid cohesion must be accompanied by a mechanism that inhibits releasing activity until cells enter mitosis. In fungi, it appears that acetylation of Smc3 by the Eco1 acetyltransferase is sufficient to block releasing activity (Rolef Ben-Shahar et al., 2008; Rowland et al., 2009; Unal et al., 2008). However, in animal cells acetylation is insufficient and stable cohesion requires binding of sororin to Pds5, an event that hinders an association between Wapl and Pds5 essential for releasing activity (Ladurner et al., 2016; Nishiyama et al., 2010; Ouyang et al., 2016).

An important question remains concerning the mechanism by which sister chromatid cohesion is generated. Specifically, what is the fate of cohesin rings during the process of DNA replication? Are the rings that entrap sisters after replication derived from rings that had entrapped individual DNAs prior to replication? In other words, do cohesin rings remain associated with chromatin during the passage of replication forks? Because Wapl-mediated turnover would mask any potential turnover induced by the passage of replication forks, we used CRISPR/Cas9 to generate a Wapl-deficient cell line in which the natural turnover of chromosomal cohesin during G1 phase is abrogated. By imaging a Halo-tagged version of Scc1 in such cells, we show that cohesin remains associated with chromatin throughout the cell cycle including S phase. We conclude that the passage of replication forks does not per se remove cohesin from chromatin.

## Results

### A pulse-chase protocol for measuring the fate of fluorescent proteins in mammalian cells

To address whether or not chromosomal cohesin is displaced by replication forks, we set out to illuminate a subset of chromosomal cohesin with a fluorescent marker in G1 and follow its fluorescence by time-lapse video microscopy as individual cells undergo S phase. To this end, we used CRISPR/Cas9 to tag all endogenous copies of *SCC1* in U2OS cells with the HaloTag (Fig. 1a). It is important to point out that Halo-tagged Scc1 was sufficiently functional to permit apparently unperturbed proliferation of the U2OS cell line. To follow the fate of nucleosome H3/H4 tetramers in the same cells, we used a transgene expressing a SNAP-tagged version of the histone variant H3.3. Transient incubation of these cells with Halo and SNAP ligands attached to different fluorescent molecules (JF549 and DY505, respectively) showed that as expected Scc1-Halo^JF549^ disappeared from chromosome arms but not centromeres when cells entered prophase (Fig. 1b), a phenomenon caused by a separase-independent release mechanism dependent on Wapl. In contrast, H3.3^DY505^ persisted throughout chromosomes during the entire cell cycle.

**Figure 1:**
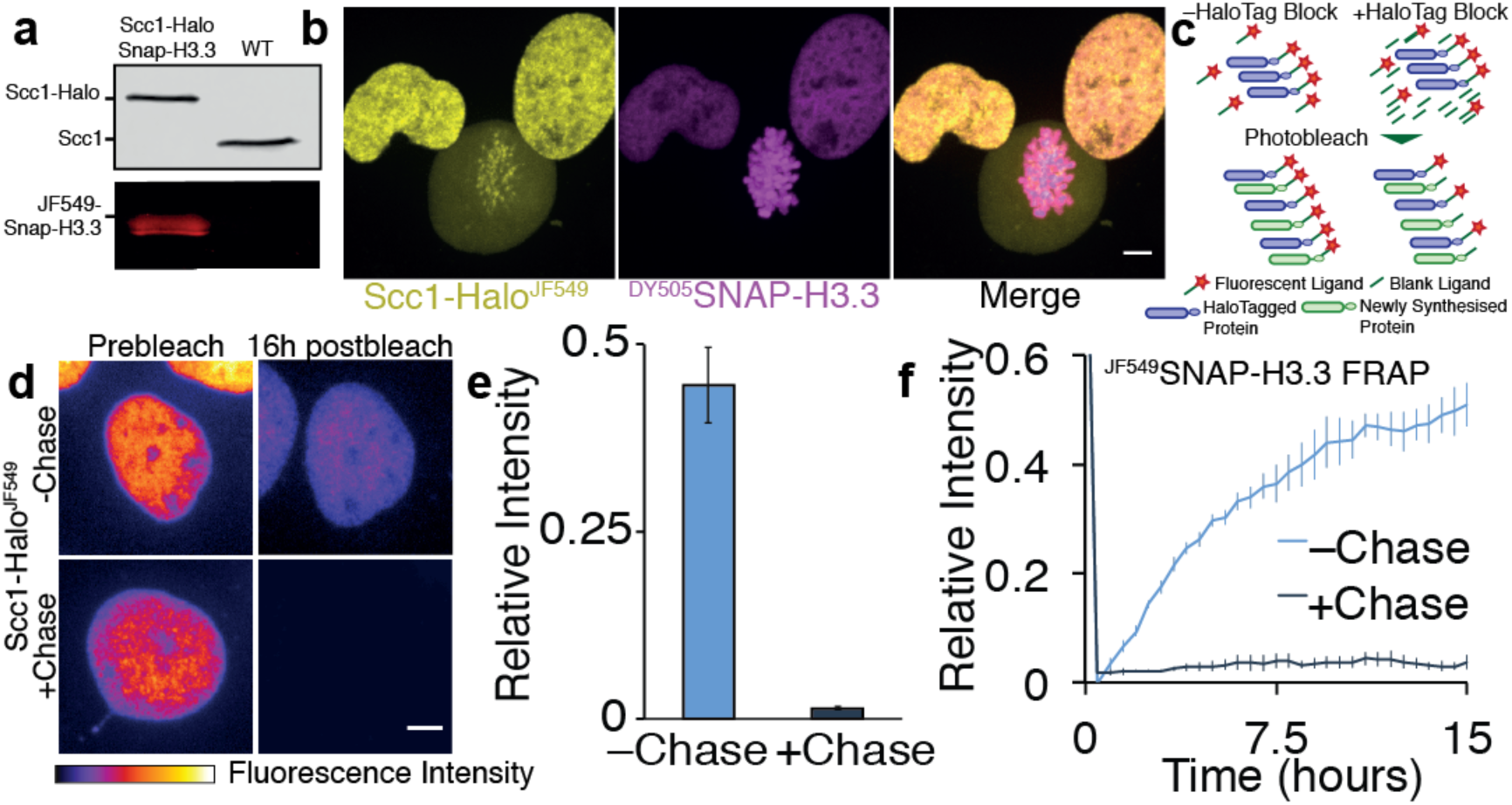
Pulse-Chase Frap (pcFRAP) permits observation of chromatin binding over long time periods. **a)** Immunoblot and In Gel Fluorescence of Scc1-Halo and SNAP-H3.3 U20S nuclear extract **b)** Live cell microscopy images of Scc1-Halo^JF549^ and ^DY505^SNAP-H3.3. Scale bar, ***5****μ*m **c)**Schematic shows how residual fluorescent HaloTag ligand labels newly synthesised HaloTag fusion proteins. Incubation with an unlabelled ligand permanently blocks new proteins from becoming labelled **d)** Average intensity projections of z-stacks from Scc1-Halo whole nuclear FRAP experiments. Scale bar, ***5*** *μ*m **e)** Mean fluorescence intensity of Scc1-Halo^JF549^ nuclei 16 hours after bleaching of whole nucleus relative to prebleach Intensity. Recovery was observed in the presence or absence of blocking HaloTag ligand. n=11 **f**) Graph depicting half nuclear FRAP for ^JF549^SNAP-H3.3. n=6.

Our goal was next to photobleach selectively a large fraction of the nucleus and follow the fate of fluorescent molecules from the unbleached part of the nucleus. However, such imaging experiments using fluorescent fusion proteins have a fundamental limitation, namely recovery of fluorescence in the bleached part of the nucleus due to fresh synthesis of the fluorescent protein or by chromophore maturation. We found that the same problem exists with Halo and SNAP tags, i.e. newly synthesized proteins interact with fluorescent ligands that remain in the medium even after repeated washes and incubations. This is not a problem when imaging for short time intervals (<10 min) but is a major problem for longer recovery periods during which significant protein synthesis occurs. Previous studies have incubated cells in the protein synthesis inhibitor cyclohexamide to mitigate this issue (Gerlich et al., 2006; Kimura and Cook, 2001; Krol et al., 2008); however, under these conditions cells cannot progress through the cell cycle and relevant interacting proteins may become depleted.

We reasoned that by adding a surplus of an unlabelled HaloTag ligand to the medium following fluorescent labelling of the HaloTag, the excess unlabelled ligand would compete with remaining fluorescent ligand for binding to newly synthesised proteins, which as a consequence would remain non-fluorescent. To test this, we incubated Scc1-Halo cells transiently with a fluorescent HaloTag ligand (100 nM), washed the cells 4 times with an intervening 30 min incubation to maximise the removal of the unbound dye, then imaged them in the presence or absence of excess unlabelled HaloTag ligand (100 *μ*M) (Fig. 1c). We then measured recovery of nuclear fluorescence following photobleaching of the entire nucleus. In the absence of unlabelled ligand, fluorescence associated with the HaloTag recovered to 45% of the pre-bleached level within 16 h. Crucially, addition of unlabelled HaloTag ligand reduced the recovery to 1% (Fig. 1d and e). We conclude that labelling of HaloTag fusion proteins and imaging them in the presence of excess unlabelled ligand makes it possible for the first time to follow defined populations of fluorescent molecules for long periods. We call this modification to the FRAP protocol pulse-chase FRAP (pcFRAP).

To demonstrate the utility of this technique, we performed half-nuclear FRAP on H3.3^JF549^, a protein thought to reside relatively stably on DNA. In the absence of SNAP-Tag inhibitor (SNAP-Cell Block), fluorescence intensity recovered to ∼50% of pre-bleach intensity within 15 h. However, if the SNAP-Tag inhibitor was added to the imaging medium, H3.3^JF549^ intensity recovered by only 4% in 15 h (Fig. 1f). In addition to validating the pcFRAP procedure, this experiment shows that there is negligible turnover of chromosomal histone H3.3. The concept behind our pcFRAP protocol is analogous to pulse-chase experiments using radioactive isotopes. We note that the fluorescence pulse-chase method has recently been used to measure protein half-lives in mammalian cells (Bodor et al., 2012; Hou et al., 2012; Huybrechts et al., 2009; Mok et al., 2013; Yamaguchi et al., 2009).

### Generation of a Wapl-deficient Scc1-Halo U2OS cell line

Wapl-dependent releasing activity causes the continual dissociation of chromosomal cohesin in G1 cells, which is balanced by de novo loading. As a consequence, the half-life of chromosomal cohesin during G1 is about 15-25 min (Gerlich et al., 2006; Hansen et al., 2016; Ladurner et al., 2016). Because of these dynamics, a subnuclear population of chromosomal cohesin marked by selective photobleaching during G1 will disappear before cells enter S phase. In other words, Wapl-mediated turnover will mask any effect of replication. To observe the latter, it is therefore essential to measure the fate of cohesin during S phase in cells in which releasing activity has been eliminated. To do this, we transfected our Scc1-Halo U2OS cell line with a plasmid expressing Cas9 and a guide RNA that induces formation of a double strand break within *WAPL’s* M1116 codon, whose mutation has previously been shown to abrogate releasing activity in HeLa cells (Ouyang et al., 2013).

Following transfection, we isolated a clonal cell line containing three different deletion alleles. Two of these changed the reading frame and are predicted to create truncated proteins, and the other one was an in-frame deletion that removed four residues, from M1116 to C1119, including E1117, which is highly conserved among animal and plant Wapl orthologs. Despite extensive screening, we failed to obtain cell lines in which all *WAPL* alleles contained frame-shift mutations, suggesting that an activity associated with *WAPL^1116-1119Δ^* is necessary to sustain the proliferation of Scc1-Halo U2OS cells. Nonetheless, cohesin turnover in G1 *WAPL^1116-1119Δ^* cells proved to be low if not entirely absent and we therefore proceeded to create two variants, one expressing SNAP-H3.3 and the other eGFP-PCNA.

Several lines of evidence suggest that separase-independent releasing activity is drastically reduced in *WAPL^1116-1119Δ^* cells. First, cohesin formed structures known as vermicelli, albeit less pronounced in this particular cell line than previously reported for quiescent mouse embryonic fibroblasts (Fig. 2b) (Tedeschi et al., 2013). This difference could be at least partly caused by the fact that unlike the *WAPLΔ* cells used by Tedeschi and colleagues our cells are dividing and as a consequence cohesin spends less time on chromatin before separase removes it. Second, most cohesin was associated with the axes of mitotic chromosomes from prophase until the onset of anaphase (Fig. 2b). This contrasts with the situation in the parental cells where the vast majority of cohesin dissociates from chromosomes when cells enter mitosis and only persists around centromeres. Thus, the Wapl-dependent prophase pathway of cohesin dissociation appears fully defective. Third, tracking of individual Scc1-Halo^JF549^ molecules revealed a major increase in the fraction with low diffusion coefficients in *WAPL^1116-1119Δ^* cells, which are bound to chromatin (Fig. 2c and d). Lastly, selective photobleaching of Scc1-Halo^JF549^ showed that unbleached areas persisted for many hours, indicating little or no turnover (Fig. 3a).

**Figure 2:**
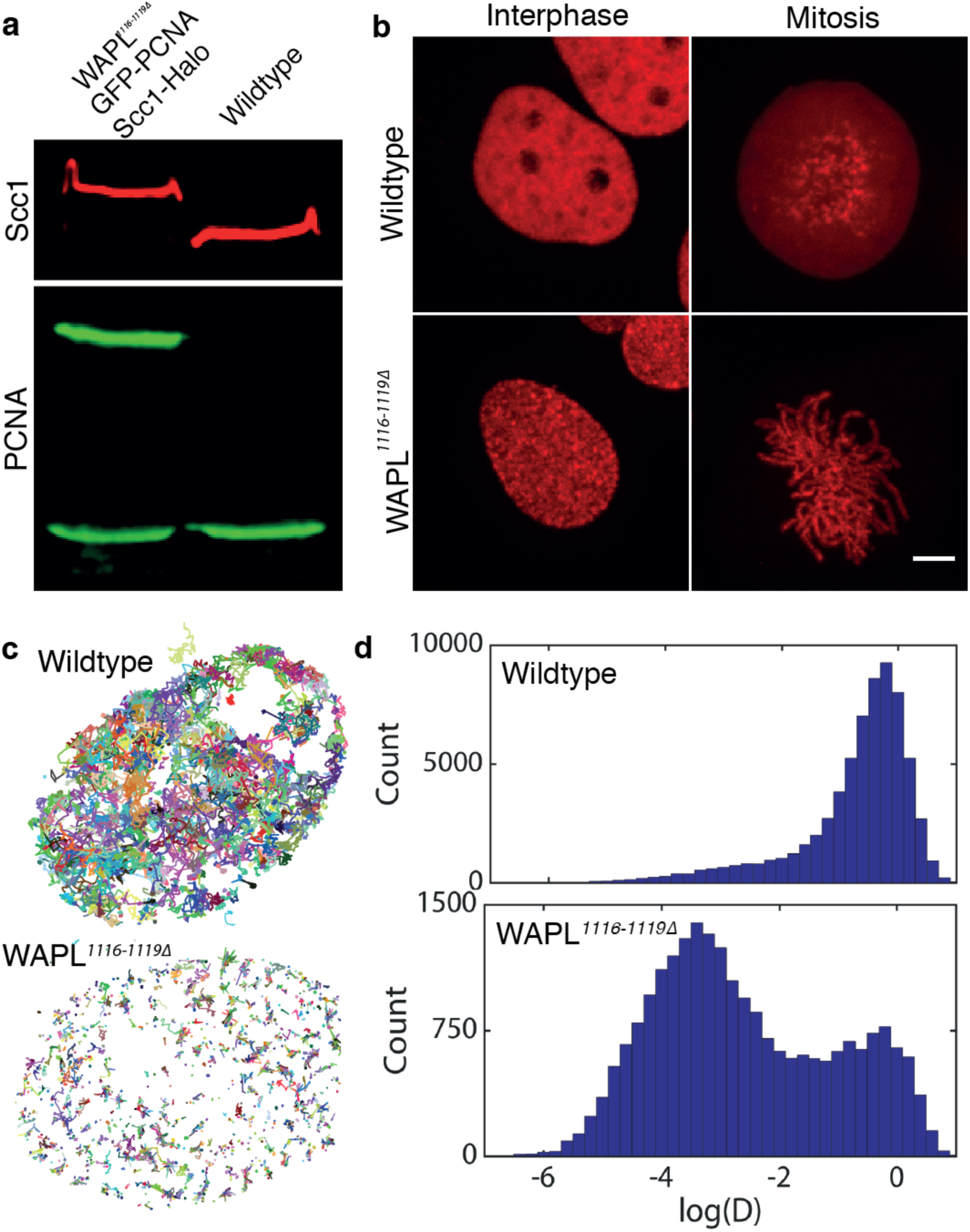
Characterisation of Sed-Halo WAPL*^1116-1119Δ^*cell line. **a)** Immunoblot of Scc1-Halo WAPL*^1116-1119Δ^* eGFP-PCNA U20S nuclear extract, **b)** Live cell microscopy images of Scc1-Halo^JF549^ in WT or WAPL*^1116-1119Δ^* U20S cells in interphase or mitosis. Scale bar, 5 *μm* **c)** Tracks of single Scc1-Halo^JF549^ molecules in wild type or WAPL*^1116-1119Δ^* cells **d)** Histograms showing the number of molecules observed moving at different diffusion coefficients

**Figure 3:**
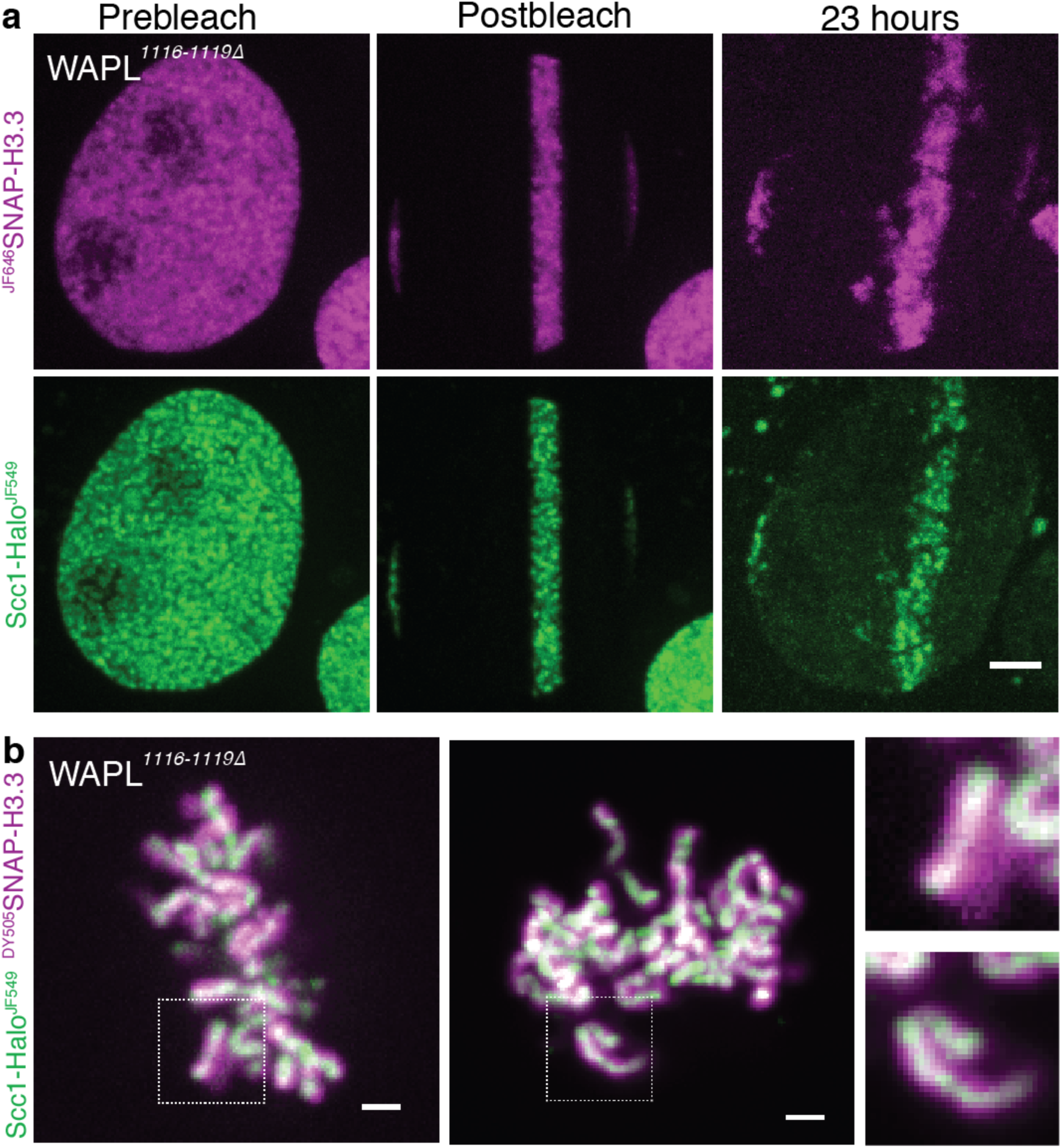
Cohesin remains associated with the same area of chromatin over long time periods. **a)** Live cell microscopy images of Scc1-Halo^JF549 JF646^SNAP-H3.3 WAPL&*^1116-1119Δ^* U20S Cells before photobleaching with a 568nm laser line, immediately after photobleaching and 23 hours later. Scale bar, 5 *μ*m **b)** Live cell microscopy images of Scc1-Halo^JF549 DY505^SNAP-H3.3 WAPL*^1116-1119Δ^* U20S cells in mitosis. Scale bar, 2 *μ*m

Another feature of *WAPL^1116-1119Δ^* cells was the low level of Scc1-Halo^JF549^ in those cells that had recently undergone mitosis. We reasoned that this might be caused by greater than normal cleavage of Scc1 by separase (Tedeschi et al., 2013). In normal cells, including our parental U2OS cell line, the cohesin that dissociates from chromosomes during prophase accumulates in the cytoplasm and is not cleaved by separase. In *WAPL^1116-1119Δ^* cells, in contrast, most cohesin remains bound to chromosomes until separase is activated at the metaphase to anaphase transition, whereupon it rapidly dissociates from chromosomes, presumably due to Scc1 cleavage. To follow the fate of cohesin as cells enter the next cell cycle, we imaged Scc1-Halo that had been labelled with JF549 during interphase and then chased with non-fluorescent ligand after cells had undergone cell division. All fluorescence associated with chromosomes disappeared during anaphase but the bulk reappeared within the nuclei of daughter cells during telophase, whereupon the fluorescence gradually decayed (Video 1). We suggest that the decay is triggered by separase cleavage during anaphase followed by degradation during the subsequent G1 period of the C-terminal Halo-tagged fragment by the N-end rule Ubr1 degradation pathway (Rao et al., 2001). In this regard, the dynamics of cohesin in *WAPL^1116-1119Δ^* cells resembles that in yeast cells, in which most cohesin is cleaved by separase due to the absence of a prophase pathway (Uhlmann et al., 1999). Because of this behaviour, daughters of *WAPL^1116-1119Δ^* cells are born with a greatly reduced pool of cohesin rings, which is only gradually replenished by de novo Scc1 synthesis.

### Lack of appreciable chromosome movement during interphase in ***WAPL^1116- 1119Δ^*** cells

Investigating through selective photobleaching experiments whether or not DNA replication per se displaces cohesin depends not only on a lack of Wapl-mediated turnover but also on limited chromosome movement as well as limited cohesin translocation. To address this, we labelled Scc1-Halo with JF549 and SNAP-H3.3 with JF646, blocked further labelling by addition of non-fluorescent HaloTag ligand and SNAP-Cell Block, photobleached all but a thin strip of chromatin within the middle of the nucleus, and recorded the fluorescence 23 hours later. The experiment was performed on cells in unknown stages of the cell cycle but most cells undergo mitosis within 30 hours of imaging. This revealed that despite movement of the cells, most fluorescence associated with both labels remained in a strip that still stretched across the nucleus (Fig. 3a). Only sporadic patches disjoined from the main strip into adjacent spaces. Crucially, fluorescence associated with the HaloTag always colocalized with that associated with the SNAP-Tag, even when the latter had moved away from the main body of fluorescence. The experiment was repeated on numerous occasions with identical results. Three conclusions can be drawn from these observations. First, neither histone H3.3 nor Scc1 dissociate in any appreciable manner from chromatin in *WAPL^1116-1119Δ^* cells over the 23 hour period of observation. Second, there is little rearrangement of chromatin throughout the nucleus during this time period consistent with previous findings (Walter et al., 2003). Third, cohesin does not translocate large distances along or between chromosomes during the time period.

Interestingly, when unbleached *WAPL^1116-1119Δ^* Scc1-Halo^JF549 DY505^SNAP-H3.3 cells entered mitosis in the absence of the prophase pathway it appeared that the bulk of cohesin was located at the intrachromatid axis (Fig. 3b). As vermicelli are evident in these cells, this suggests that much of the cohesin in *WAPL^1116-1119Δ^* cells is entrapping sister DNAs as well as holding loops. Whether this is really the case and if so how is an interesting question.

### Most cohesin remains within local chromatin zones throughout S phase

Though the previous experiment demonstrated Scc1’s stable association with specific zones of chromatin over long periods in *WAPL^1116-1119Δ^* cells, it did not directly address cohesin’s fate during DNA replication. To do this, we repeated the experiment using *WAPL^1116-1119Δ^* cells expressing eGFP-PCNA instead of SNAP-H3.3. Our goal was to label a restricted zone of Scc1-Halo during G1, image eGFP-PCNA sufficiently frequently to establish passage through S phase, and then record the pattern of Scc1-Halo fluorescence once S phase had been completed. Cells were determined to be in S phase depending on the pattern of eGFP-PCNA, which accumulates transiently in patches of replicating chromatin only during DNA replication.

Due to cleavage of most Scc1 by separase in *WAPL^1116-1119Δ^* cells during anaphase, fluorescence associated with G1 cells was much lower than in S phase or G2 cells, with the result that Scc1-Halo^JF549^ images were fainter than would otherwise have been the case. Nevertheless, we were able to record defined segments of Scc1-Halo fluorescence in numerous cells (n=10) before and after cells had unambiguously completed S phase. In all cases, the subnuclear pattern of Scc1-Halo^JF549^ fluorescence in G2 cells resembled that of their G1 precursors (Fig. 4). We conclude that the passage of replication forks does not displace cohesin from chromatin in a manner that would cause it to diffuse appreciably within the nucleus before reloading. Inevitably, our experiment does not exclude the possibility that in *WAPL*+ cells, replication forks cause dissociation by inducing Wapl-dependent releasing activity. The key point is that our observations demonstrate that cohesin in fact can persist on chromatin throughout replication, at least under conditions in which it cannot be removed by Wapl-mediated releasing activity.

**Figure 4:**
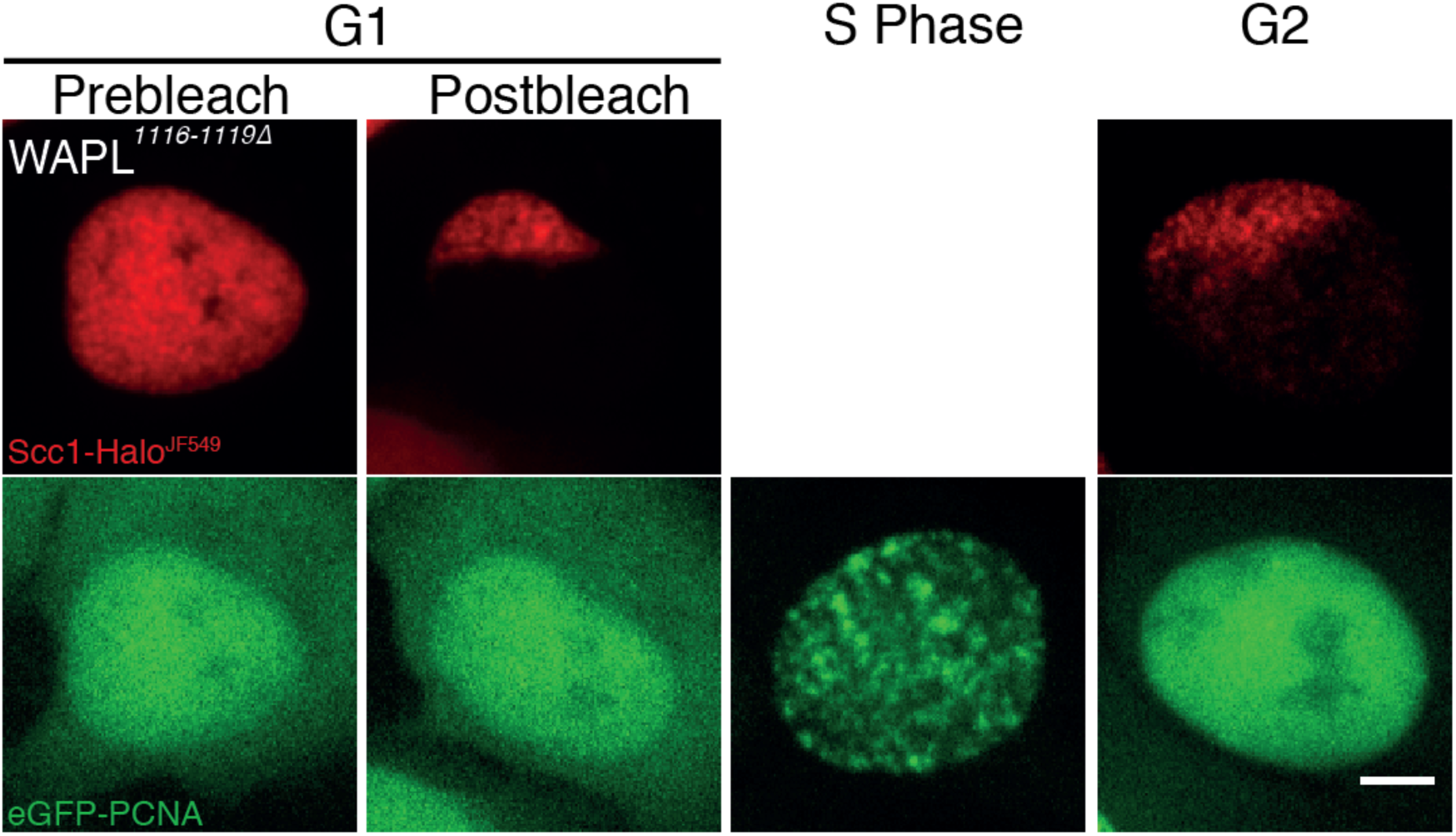
A pool of cohesin remains bound to DNA during S-phase. Live cell microscopy images of SCC1-Halo^JF549^ WAPL*^1116-1119Δ^*eGFP-PCNA cells in G1 before photobleaching, in G1 after photobleaching, in S-Phase and in G2. It was not possible to image SCC1-Halo^JF549^ between G1 and G2 because of low starting signal in G1. This fluorescence was lost easily by photobleaching from eGFP and JF549 acquisition. Scale bar, 5 *μ*m

## Discussion

This paper set out to address a key question concerning the dynamics of cohesin on chromosomes, namely whether cohesin rings associated with chromatin fibres during G1 are necessarily displaced by the passage of replication forks. Answering this question is important with regard to the mechanism by which cohesion between sister DNAs is generated during S phase. If cohesin rings entrap individual DNAs during G1 and a pair of sister DNAs during G2, then the latter could in principle be derived from the former by extrusion of forks through cohesin rings. If so, cohesin rings should not be displaced by the passage of replication forks.

We therefore set out to determine whether displacement of cohesin from chromatin is an obligatory aspect of replication fork progression. Using CRISPR/Cas9 to create a U2OS human cell line whose Scc1 is tagged with the HaloTag and that lacks releasing activity due to deletions within *WAPL*, we have been able to image subnuclear zones of JF549-labelled cohesin throughout the cell cycle and show that they persist throughout S phase. This proves that replication forks do not cause cohesin to dissociate from chromatin in a manner that permits its diffusion throughout the nucleus before re-associating.

Due to the low resolution of our imaging, we cannot exclude the possibility that cohesin dissociation does in fact take place locally during replication fork progression but that it re-associates with chromatin so rapidly that it does not leave the unbleached region. However, in mouse ES cells the time between cohesin release and loading events is ∼33 minutes (Hansen et al., 2016), therefore a replication specific pathway would be needed to explain such a high re-association rate. To address this possibility it would be necessary to image individual cohesin molecules during passage of replication forks, which is currently not technically possible in living cells due to their size. Such an experiment might be feasible in Xenopus extracts where it is possible to use TIRF microscopy to image cohesin during DNA replication. Preliminary observations of this nature have recently been reported but the data are not conclusive (Kanke et al., 2016). Importantly, had we found that chromosomal cohesin was recycled throughout the nucleus following passage of replication forks then we would know that DNA replication displaces cohesin. Our findings to the contrary demonstrate that cohesin like the histone H3/H4 tetramer has the ability to persist on chromatin during replication fork passage.

## Experimental Procedures

### Antibodies

The following commercial antibodies were used: Scc1 (ab154769, Abcam), PCNA (ab29, Abcam).

### HaloTag/SNAP-Tag Ligands

JF549-SNAPTag, JF646-SNAPTag, JF549-HaloTag, JF646-HaloTag were as previously described (Grimm et al., 2015). DY505-SNAPTag and SNAP-Cell Block were purchased from NEB, UK. The blocking ligand for pulse-chase FRAP was the HaloTag^®^ succinimidyl ester (O2) ligand (Promega). The reactive group was inactivated by incubating the ligand with 1 M Tris–HCl (pH 8.0) for 1 h at 25°C as previously described (Yamaguchi et al., 2009). The HaloTag amine ligand was tested but had no effect, possibly due to low cell permeability.

### Plasmids

pSpCas9(BB)-2A-Puro (PX459) V2.0 was a gift from Feng Zhang (Addgene plasmid # 62988). Human H3.3 cDNA was a gift from Danette Daniels (Promega). pEGFP-PCNA-IRES-puro2b was a gift from Daniel Gerlich (Addgene plasmid # 26461) and was used to generate pEGFP-PCNA-IRES-BSD and pSNAPTag-H3.3-IRES-BSD. Scc1-HaloTag HR template (1kb homology arms) was cloned into pUC19 between KpnI and SalI.

### Guide RNAs

The following guide RNAs were inserted into the BbsI restriction site of pSpCas9(BB)-2A-Puro (PX459) V2.0 (Ran et al., 2013).

SCC1 C CCAAGGTTCCATATTATATA

WAPL M1116 GCATGCCGGCAAACACATGG

### Cell Lines

Scc1-Halo U2OS cells were generated by cotransfection of pX459_SCC1 C and the Scc1-HaloTag HR template. Cas9 expressing cells were selected with puromycin (2 *μ*g/ml) for 2 days and plated for colony picking. Homozygous clones were identified by PCR and confirmed by Western blotting. The Scc1-Halo, SNAP-H3.3 U2OS cell line was generated by stable integration of pSNAPTag-H3.3-IRES-BSD into the Scc1-Halo cell line. To generate the Scc1-Halo, *WAPL^1116-1119Δ^*, eGFP-PCNA or SNAP-H3.3 cell line pX459 WAPL M1116 was transiently expressed in Scc1-Halo U2OS cells and pEGFP-PCNA-IRES-BSD or pSNAPTag-H3.3-IRES-BSD was stably expressed. Clonal stable cell lines were isolated using blasticidin (5 *μ*g/ml) selection.

### Conventional and pulse-chase labelling

One day before imaging, U2OS cells were seeded on glycine coated glass bottom dishes. Glycine coating of coverslips for 15 min at room temperature greatly reduces binding of TMR and JF549 to glass(van de Linde et al., 2011). Cells were then incubated with fluorescent HaloTag ligands JF549 and JF646 (100 nM) for 15 min and SNAP-Tag ligands JF549 (100 nM) and DY505 (1 *μ*M) for 30 min. Cells were washed in CO_2_ equilibrated medium 3 times, then incubated for 30 min to allow the ligand to exit the cells. The medium was replaced twice more for a total of 5 washes.

For pulse-chase FRAP using the SNAP-Tag, SNAP-Cell^®^ Block was added to a final concentration of 10 *μ*M. Halo blocking ligand was used at 100 *μ*M.

### Microscopy

Live-cell imaging was performed on a spinning disk confocal system (PerkinElmer UltraVIEW) with an EMCCD (Hamamatsu) mounted on an Olympus IX8 microscope with an Olympus 60× 1.4 N.A objective. A custom single-molecule fluorescence microscope with a 561 nm (Oxxius) excitation laser was used to record PALM movies on an EMCCD camera (Andor) at 15.26 ms/frame for 20,000 frames.

### PALM Analysis

Data analysis and simulations were performed in MATLAB (MathWorks). PSFs were localized to 40 nm precision by elliptical Gaussian fitting. Localisations within a radius of 0.57 μm in consecutive frames were linked to tracks. Tracks with more than four steps were used to compute apparent diffusion coefficients (*D\**).

## Contributions

KN and JR conceived the study and designed all experiments. JR performed all experiments. JH and BR shared unpublished results and helped with the inactivation of Wapl. JG and LL synthesised Janelia Fluor dyes.

## Acknowledgments

We thank Danette Daniels for allowing us to test Promega’s HaloTag building blocks for pcFRAP and for providing the H3.3 cDNA. We are grateful to Stephan Uphoff for helping with the single molecule imaging. We are also grateful to Bungo Akiyoshi, Anders Hansen and members of the Nasmyth group for comments on the manuscript. We thank Micron for maintaining the microscopes.

